# Incorporating extrinsic noise into mechanistic modelling of single-cell transcriptomics

**DOI:** 10.1101/2023.09.30.560282

**Authors:** Kaan Öcal

**Affiliations:** School of Informatics, University of Edinburgh, Edinburgh EH8 9AB, UK; School of Biological Sciences, University of Edinburgh, Edinburgh EH9 3JH, UK

## Abstract

A mechanistic understanding of single-cell transcriptomics data requires differentiating between intrinsic, extrinsic and technical noise, but an abundance of the latter often obscures underlying biological patterns. Accurately modelling such data in the presence of large cell-to-cell heterogeneity due to factors such as cell size and cell cycle stage is a challenging task. We propose a tractable, fully Bayesian framework for mechanistic modelling of single-cell RNA sequencing data in the presence of cellular heterogeneity. Applied to murine transcriptomics data, we show that cell-specific effects can significantly alter previously inferred dynamics of individual genes. Our implementation is statistically exact and readily extensible, and we demonstrate how it can be combined with Bayesian model selection to compare various models of gene expression and measurement noise.

## 1 Introduction

Single-cell RNA sequencing (scRNAseq) allows us to observe and measure gene expression in thousands of genes and cells at a time, with a wide range of applications in fields such as immunology, cancer research and developmental biology. While large-scale analyses of expression patterns across cells and tissues have been in the focus for the past several years, the application of scRNAseq to understand gene expression on a mechanistic level has been a more recent concern. Two major reasons for this are the substantial technical noise encountered in most scRNAseq methods [1, 2], as well as the limited type of data obtained by classical scRNAseq experiments, resulting in frequent parameter unidentifiability [3, 4].

Technical noise in scRNAseq can arise from various factors such as dropout [5], phenotypic heterogeneity in tissues, and insufficient read depth and gene coverage [6, 7]. To mitigate the low signal-to-noise ratio in raw scRNAseq data, experimental reads are typically subjected to extensive quality control, normalisation and clustering steps [1] before further analysis. While such methods are well established for investigating global mRNA transcription and differential gene expression patterns, their reliability for mechanistic analyses of gene expression has been less investigated. On a biological level, transcription is strongly influenced by various cell-specific factors such as cell size, cell cycle stage and cell type. The latter two can often be discerned based on observed transcription patterns, and cell size remains as the largest contributor of extrinsic noise for most genes [8], frequently correlating with global transcription levels to induce concentration homeostasis [9, 10].

In this paper we propose a fully Bayesian framework for the mechanistic analysis of scRNAseq data based on the telegraph model of gene expression [11], building on [3, 12, 13] and including cell size effects via a cell-specific scaling factor. Our framework is implemented in the probabilistic programming language Stan [14] and relies on a state-of-the-art Hamiltonian Monte Carlo sampler. We investigate mechanistic explanations for individual gene expression levels, and use Bayesian model selection to distinguish between modes of gene expression and evaluate the possible presence of zero-inflation. Following [13], our approach does not rely on preprocessing steps such as normalisation or imputation [15].

We test our model on an allele-specific dataset of mouse fibroblasts taken from [4], as well as a subset of the (non-allele-specific) mouse embryonic stem cell (mESC) dataset in [16]. While the standard telegraph model provides a good empirical fit to mRNAseq data on the level of marginal distributions, our results suggest that this agreement may be less fundamental than commonly thought. Indeed, taking cell sizes into account we reinterpret experimental mRNA distributions as mixtures of heterogeneous distributions based on cell-specific factors. This reinterpretation provides an alternative mechanistic explanation of the data, but it also predicts significantly different parameters from the classical telegraph model: accounting for cell-specific effects affects burst size, frequency and duration on a global level, in accordance with recent observations [17, 18]. While we exclusively focus on cell-size effects in this paper, our proposed framework is easily extended to include further sources of heterogeneity such as cell cycle stage and cell-specific capture efficiencies [19], depending on the type of data available.

## 2 Methods

### Mechanistic models of gene expression

The telegraph model of gene expression describes a gene with two possible states (*G*_on_ and *G*_off_) and mRNA *M*, together with the following set of reactions:

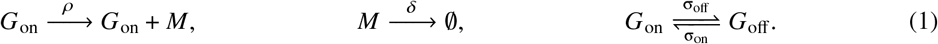

We can normalise the rates such that δ = 1 without changing the steady-state distribution (mRNA degradation rates can often be measured directly [20]). The steady-state distribution over mRNA numbers predicted by the Chemical Master Equation can be written in terms of the confluent hypergeometric function _1_*F*_1_ as follows [11]:

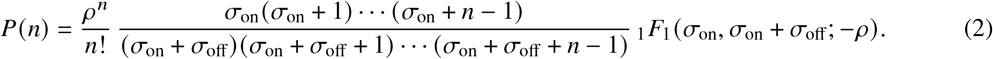

A ubiquitous simplification of the telegraph model is obtained in the bursty limit where σ_off_, ρ→∞, with ρ/σ_off_ =: *b* constant (equal to the mean burst size). The steady-state distribution for this bursty model is a negative binomial distribution, which is commonly used to model scRNAseq data:

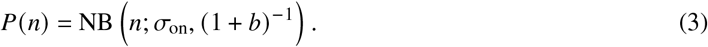

If we denote by 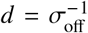 the average time the gene spends in the on state, the bursty limit can be seen as a simplification of the telegraph model where *d*≈0.

The constitutive, or Poisson model of gene expression is obtained from the bursty model in the limit *b*→0 and σ_on_ →∞with *b*σ_on_ =: λ constant, equal to the production rate. Here the steady-state distribution equals Poi(λ). Since the Fano factor of (3) equals 1+*b*, while that of a Poisson distribution is exactly 1, bursty gene expression is often used to model experimental overdispersion of mRNA counts in prokaryotic and mammalian scRNAseq data. This can be misleading in the presence of extrinsic noise, which often results in inflated variance estimates. For example, the constitutive model with a Gamma-distributed transcription rate also predicts a negative binomial steady-state distribution [21], and we will see that putative high values of *b* can often be explained by such cell-specific scaling factors.

Certain parameter combinations for the telegraph model can be practically unidentifiable from steady-state data due to the existence of the bursty and constitutive limits. In the bursty regime, the steady-state distribution resembles (3), which only depends on σ_on_ and the ratio ρ/σ_off_, and it becomes difficult to identify ρ and σ_off_ separately. Similarly, near the constitutive limit the steady state distribution approaches Poi (*b*σ_on_), and *b* and σ_on_ become separately unidentifiable. Inferring the parameters individually requires the use of time-course data, which unfortunately is not commonly available with scRNAseq methods.

Since gene expression in mammalian cells often approaches the bursty regime, this issue is noticeable in practice. Arbitrarily large values of ρ and σ_off_ explain the data to a similar degree, and obtaining reasonable parameter estimates requires placing strong priors on parameters, which in turn heavily biases inference results. Even for very large sample sizes, maximum likelihood estimation will suffer from large uncertainty for both parameters, as measured by the Fisher information matrix or profile likelihoods approaches. For this reason, [4] focusses on identifying specific “summary statistics”, such as the mean burst size and bursty frequency, that are more readily identifiable and give insight into gene expression dynamics.

Continuing this line of reasoning, we reparametrise the telegraph model using the following three parameters:

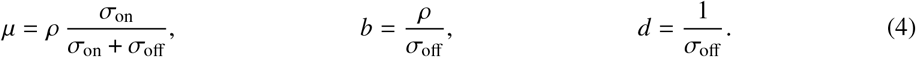

Here *μ* is the mean number of mRNAs, *b* is the average number of mRNA produced per burst (consistent with the standard definition in the bursty limit), and *d* is the average burst duration, normalised by the average mRNA lifetime. This approach exhibits the bursty and constitutive models as special cases of the telegraph model rather than as limits: the bursty model is obtained when *d* ≈ 0, and the constitutive model when additionally *b*≈0. This reparametrisation bypasses the structural identifiability issues pointed out above, and the question of which model to use — telegraph, bursty or constitutive — reduces to testing the hypothesis that *d*≈0 or *b, d*≈0, which can be done using Bayesian model selection. The mechanistic parameters can be recovered by inverting (4), as shown in SI A.

### Technical noise

A common source of technical noise in scRNAseq data is the low capture efficiency β of many methods [6, 7]. Assuming that every mRNA molecule is captured with probability β independently of the others, we observe a binomial thinning of the true mRNA numbers. It can be shown (see SI A) that accounting for global capture efficiency in this way simply scales ρ by β. As a result, ρ and β are jointly unidentifiable, and separating the two requires estimation of β. In our parametrisation, both μ and *b* are scaled by β and have to be corrected for β < 1. Since spike-in data is not commonly available for UMIs [22], we are currently unable to perform this correction, but our results are mostly independent of β.

Zero-inflation [23] is another source of technical noise for many protocols. To model possible zero inflation we include an additional parameter *p*_0_ ∈[0, 1] that controls the probability that a read will return 0 mRNA due to technical dropout (see SI A for details). We can then compare the model evidence for the original and the zero-inflated model by computing Bayes factors.

As most scRNAseq datasets are not allele-specific, the observed mRNA counts *n* = *n*_1_+*n*_2_ typically comprise reads from two alleles (in diploid cells, ignoring gene replication). Assuming that both alleles are expressed independently, we can model this as the sum of two independent telegraph models. We will assume that both alleles have the same kinetic parameters; an earlier analysis [24] of mouse blastocyst data detected no significant allele-specific effects in the majority of genes.

### Cell size dependence

Total mRNA expression levels reported by scRNAseq differ widely between cells, even among cells of the same type. This can be due to differences in capture efficiency between cells [19], or due to biological factors such as cell volume and the cell cycle. In this paper we will concentrate on cell size effects to complement recent research [25, 26] on analysing cell cycle stages. To model (partial) concentration homeostasis we introduce a dependence between cell volume and transcription rates in the shape of

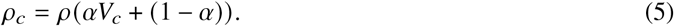

Here *V*_*c*_ is a cell-specific volume factor, shared between all genes, and α∈[0, 1] captures the volume dependence of the gene (α = 0 implies no volume dependence, while α = 1 represents full concentration homeostasis). (5) is based on the observation in [8] that not all genes scale equally with cell size; we will refer to this extension of the telegraph model as the extrinsic noise model. Eq. (5) assumes that cell volume modulates the average burst size *b*, as opposed to the burst frequency σ_on_, consistent with [9]; a more detailed analysis could additionally include cell size effects on these parameters as in [18].

As we do not have access to absolute cell volumes, we normalise the *V*_*c*_ to have mean 1 and focus on relative differences in expression. The factors *V*_*c*_ can be interpreted as cell-specific corrections to the global transcriptional volume, which can be affected by more than just the physical cell volume. However, in [8] cell size was reported to be the biggest source of extrinsic noise for most genes, and the observed strong correlation between cell size and global mRNA levels suggests that the *V*_*c*_ are a useful proxy for physical cell size.

Incorporating cell-size effects allows us to perform a variance decomposition of the form

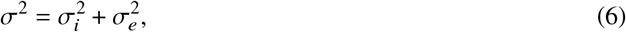

where 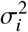 represents intrinsic noise from the telegraph model and 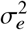represents extrinsic noise due to differences in the *V*_*c*_. The fraction of extrinsic noise for each gene can be estimated by linearly regressing mRNA counts against the volume factors *V*_*c*_, or from the model (SI E). We verify that both approaches yield similar predictions, and that the fraction of extrinsic noise is consistent across alleles and experimental replicates.

### Bayesian inference

A fully Bayesian mechanistic analysis of scRNAseq datasets is computationally challenging [19]. While the telegraph model is relatively simple, standard inference approaches do not scale well to computing posteriors over hundreds of genes with hundreds of observations each. In particular, the steady-state solution given by (2) involves confluent hypergeometric functions that are hard to evaluate quickly and accurately, and suffer from numerical issues in the (very common) bursty regime. Following [3] we therefore use an augmented MCMC sampler [27] based on an alternative representation of the steady-state telegraph model, the Beta-Poisson model derived in [28] (see SI A). As Gibbs sampling for a simplification of this model was observed in [13] to mix slowly, we used Hamiltonian Monte Carlo (implemented in Stan [14]) to explore the posterior. This allows us to consider both the telegraph model, which describes one allele, and the combination of two independent telegraph models to describe pooled allele data. The models we implemented are described in SI A, and the Stan source code is made available with this paper.

For the standard telegraph model, parameters can be inferred separately for each gene. In contrast, including the cell-specific parameters *V*_*c*_ induces a coupling between genes, and the posterior has to be computed jointly for all genes. As this is computationally very demanding, we perform inference in batches of several dozen genes each; we will see that our estimates of *V*_*c*_ are highly consistent across batches. In practice it is sufficient to estimate these parameters for each cell based on one batch of about 50 genes, and to fit the remaining genes separately based on these estimates.

## 3 Results

Our first analysis was performed on data from 224 primary adult fibroblasts, predominantly in the G1 stage, published in [4] and containing data for two alleles, CAST/EiJ and C57BL/6. To reduce the effects of technical noise we excluded genes with mean expression < 10 for both alleles. The resulting 459 genes were split into 9 equal batches for the extrinsic noise analysis.

Our second dataset consisted of mouse embryonic stem cells (mESCs) published in [16]. To reduce celltype noise we considered those cells identified as belonging to a randomly chosen cluster (cluster 9) in [16]. We truncated each of the three sequencing replicates to 460 cells, the minimum across the three, and again excluded genes with mean expression < 10 for all replicates. The resulting 262 genes were split into 5 batches of size 44 and one of size 42 for the extrinsic noise analysis.

Both the telegraph model and the extrinsic noise model reproduce experimental mRNA distributions, but with markedly different dynamics (Figs. 1, S1 and S2). While the standard telegraph model typically predicts clearly bursty dynamics characterised by *d* ≈0, this is not the case for the extrinsic noise model (cf. [17]). Based on the extrinsic noise model we can identify distinct groups of cells depending on the volume factors *V*_*c*_, and we can separate the observed mRNA distributions into contributions from these groups. The extrinsic noise model thus recovers a more detailed picture of transcription across cells that provides an alternative mechanistic explanation of the data. Estimates of the volume factors are highly consistent (ρ > 0.95) across batches (Figs. S4 and S6), and can be reliably inferred from a few dozen high-expression genes.

**Figure 1:**
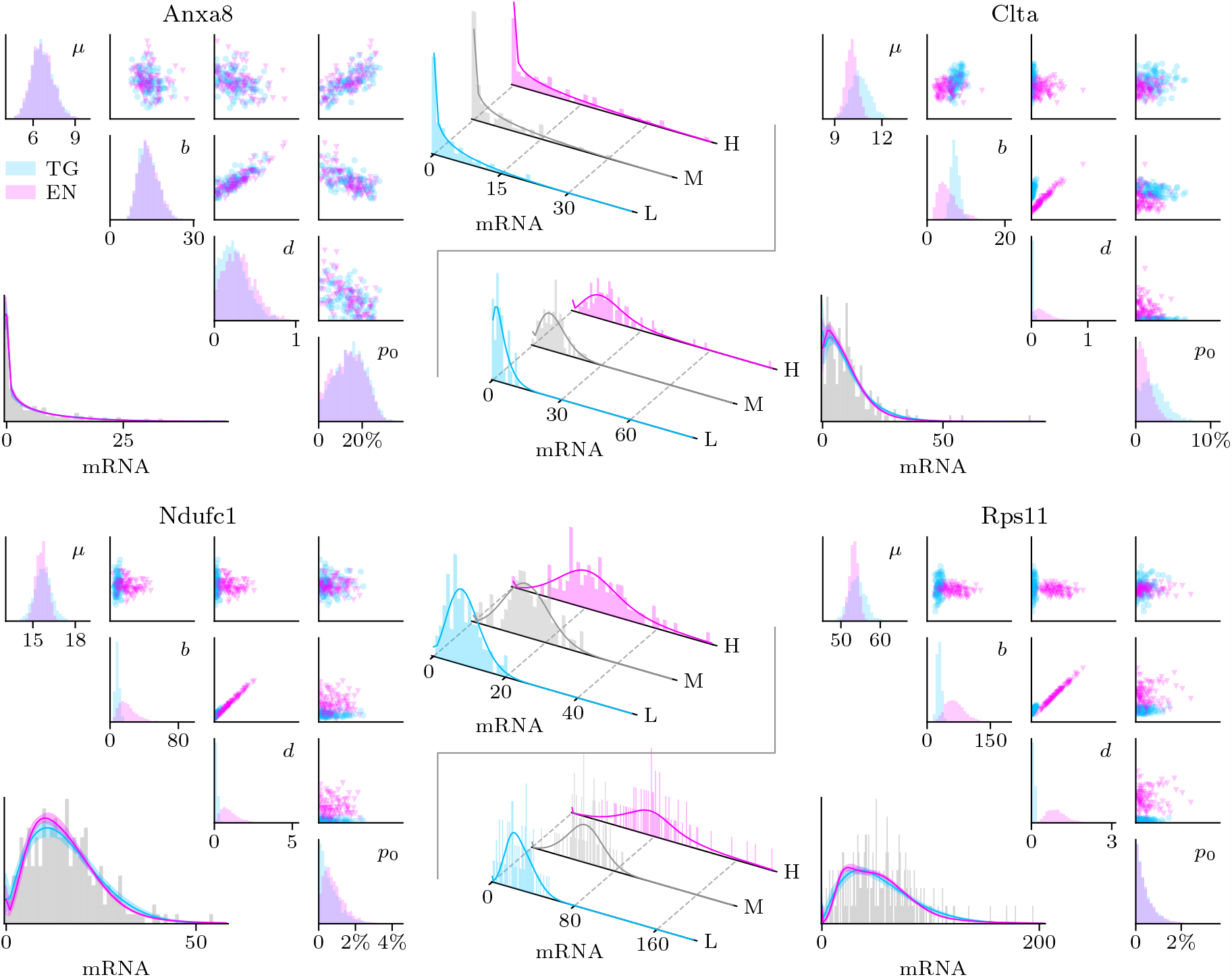
Inference results for the telegraph and extrinsic noise models, showing 4 selected genes (C57 allele). The triangular corner plots show posterior marginals (diagonal) and pairwise scatterplots (above diagonal). The histograms below the diagonal show the observed mRNA distributions, together with posterior predictive distributions for both models (solid lines represent posterior means, shaded areas 90% credible intervals). Both models reproduce experimental distributions with high confidence, despite significant differences and uncertainty in parameters. The center plots decompose observed mRNA distributions into contributions from low (L), mid (M) and high-expression (H) cells for the extrinsic noise model (the classification was performed by splitting the cells, sorted by *V*_*c*_, into three groups of equal size).

Accounting for extrinsic noise changes estimated transcription dynamics genome-wide. The telegraph model consistently underestimates the burst duration *d* compared to the extrinsic noise model (Figs. 2, 3 and S3), and the on rate σ_on_ exhibits a similar pattern. The extrinsic noise model thus predicts genes to spend more time in the on state, as was previously suggested in [17]. For genes with *d*≈0, the telegraph model falsely interprets the extrinsic noise as being attributable to larger burst sizes *b*; interestingly, this trend is reversed for genes that are predicted to be far from the bursty limit.

**Figure 2:**
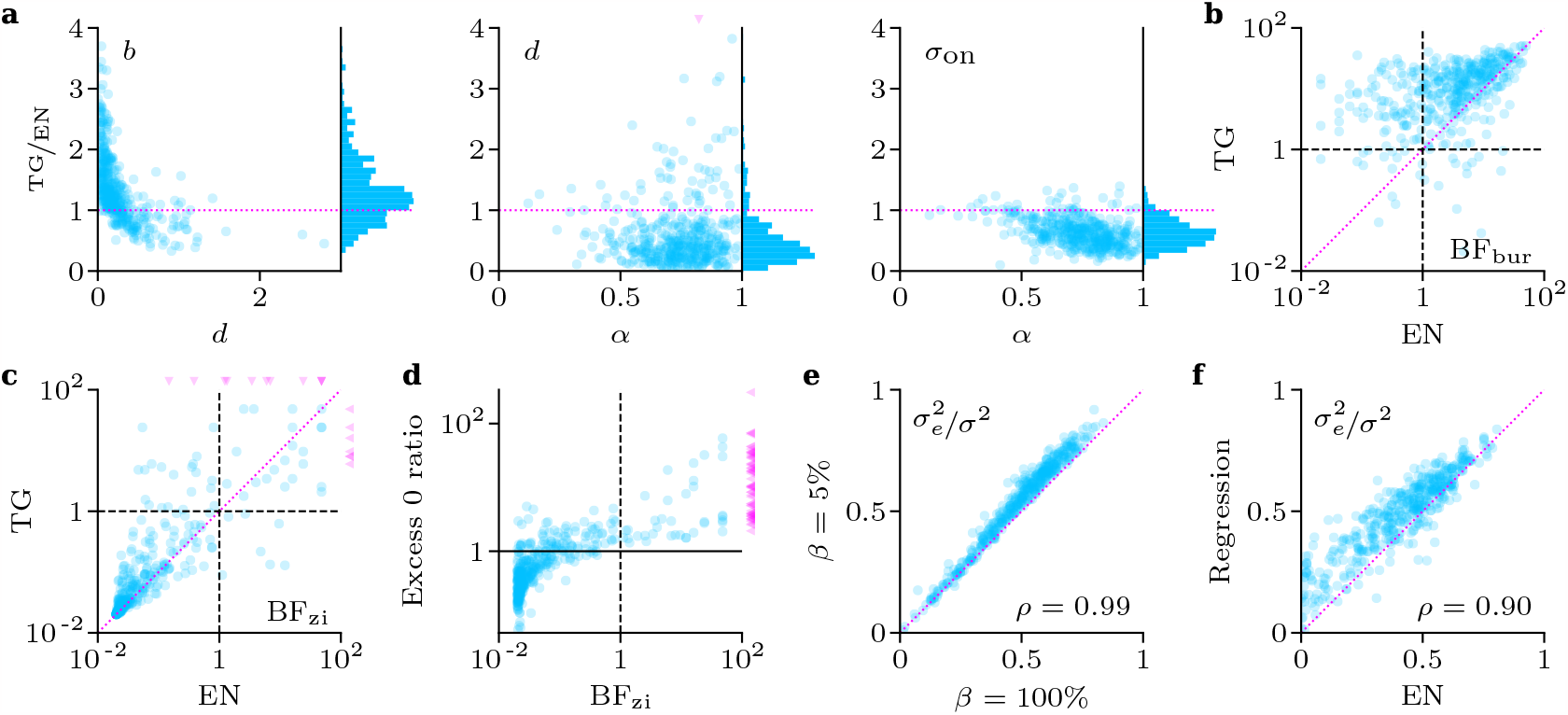
Analysis of the single-allele mouse fibroblast dataset (C57 allele) using the telegraph (TG) and extrinsic noise (EN) models. **a)** Comparison of parameter estimates under both models, obtained as posterior means. Shown are the ratios of inferred parameters against the burst duration *d* and volume scaling factor α estimated for the EN model. **b)** Including extrinsic noise results in a systematic decrease of Bayesian evidence for the bursty model of gene expression, as measured by the Bayes factor. **c)** Evidence for zero inflation is consistent between the two models, indicating its presence in a minority of genes. Triangles indicate values outside axis limits. **d)** Comparing observed and predicted abundances of zero counts measures evidence in favour of zero inflation (see SI D). **e)** Estimates of the fraction of extrinsic noise are not strongly affected by the global capture efficiency β (see SI E). **f)** The fraction of extrinsic noise predicted by the EN model agrees with that obtained by regressing mRNA counts against volume factors.

**Figure 3:**
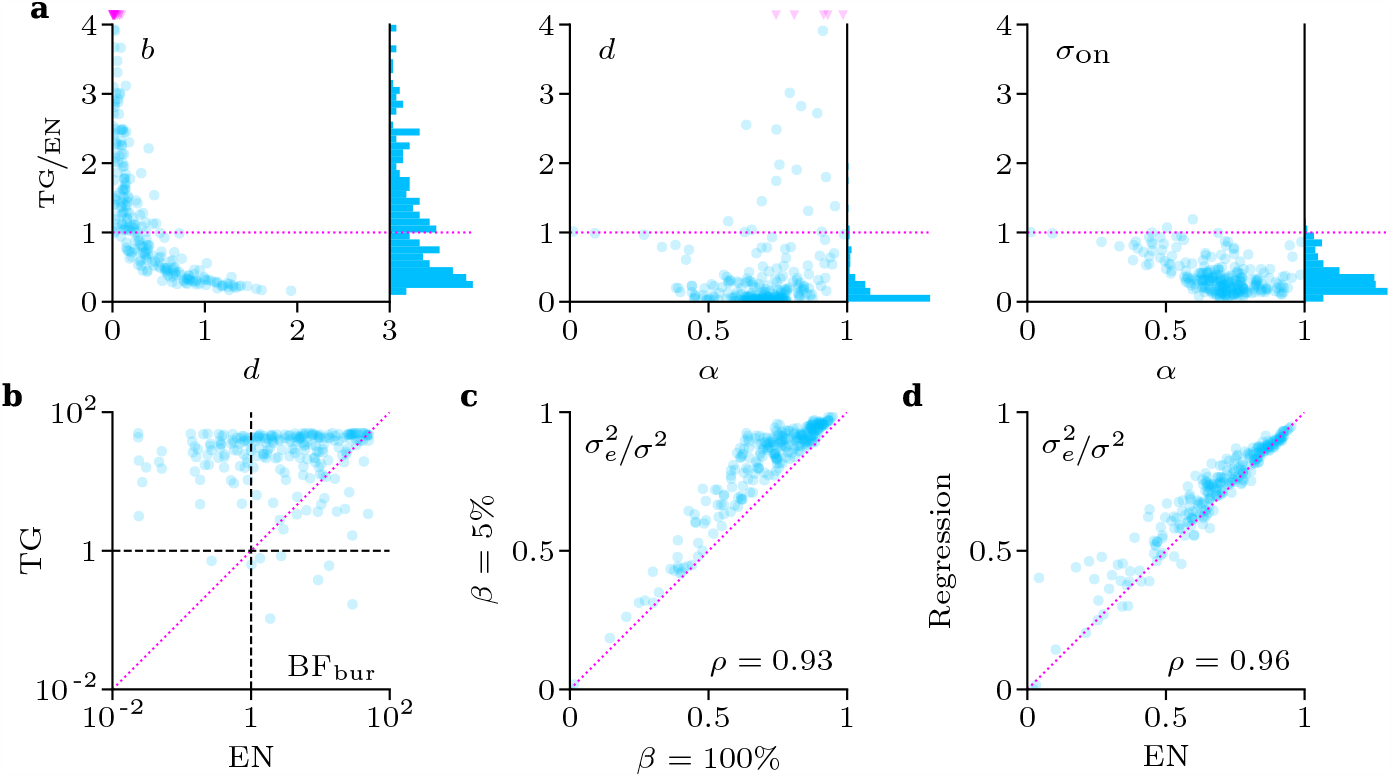
Analysis of the two-allele mESC dataset (replicate 1) using the telegraph (TG) and extrinsic noise (EN) models. The parameters μ and *b* are normalised for one allele as described in SI A. **a)** Comparison of parameter estimates under both models, obtained as posterior means. Shown are the ratios of inferred parameters against the burst duration *d* and volume scaling factor α estimated for the EN model. **b)** Including extrinsic noise results in a systematic decrease of Bayesian evidence for the bursty model of gene expression. **c)** Estimates of the fraction of extrinsic noise are not strongly affected by the global capture efficiency β . **d)** The fraction of extrinsic noise predicted by the EN model agrees with that obtained by linear regression of mRNA counts against volume factors.

For the single-allele fibroblast dataset we find evidence in favour of zero inflation in a minority of genes, irrespective of the model used (Figs. 2 and S3). The extent of zero inflation, however, is typically small. The mESC dataset shows neglible amounts of zero inflation throughout, as does the pooled fibroblast dataset discussed below.

Estimates of the fraction of extrinsic noise under our model were remarkably robust to the global capture efficiency β (Figs. 2, 3 and S3), and were consistent with a linear regression estimate against the volume factors *V*_*c*_. In agreement with [13], we found that extrinsic noise accounted for 50-70% of the variance across many genes, keeping in mind that this estimate will likely be lower for genes with lower expression levels.

We next compared results for the single-allele fibroblast data with those obtained from pooling the alleles, using the pooled extrinsic noise model for the latter. We found that estimates generally agreed between alleles, and between single-allele and joint datasets (Fig. S5), but notably *p*_0_ was generally negligible in the joint case, with the exception of the gene *thy1* involved in fibroblast differentiation [29]. This mirrors our results for the mESC dataset; it is reasonable that zero-inflation for pooled alleles is only detected if both alleles drop out simultaneously, which is significantly rarer than one allele dropping out. The very high consistency of our volume factor estimates *V*_*c*_ suggests that our extrinsic noise estimates are not affected by allele-specific differences.

For the mESC dataset, which is not allele-specific, we consistently found high correlations (ρ > 0.8) between parameters inferred using the telegraph model and its two-allele version (not shown). We finally compared the inferred parameters across the replicates of the mESC dataset (Fig. S7). Mean expression μ correlated very strongly across replicates, with the mean burst size and duration correlating more weakly, which suggests that the inferred parameters are somewhat susceptible to batch effects. Notably, this was not the case for the volume dependence parameter *α*, although the estimated fraction of extrinsic noise remained quite similar between replicates. This somewhat mystifying observation suggests that the parameter *α*, while useful in making predictions, may not in itself be a physically robust quantity, a point that will merit further research.

## 4 Discussion

In this paper we propose a fully Bayesian framework, similar in vein to [3, 13, 19], for investigating the effect of extrinsic noise on mRNA counts obtained using single-cell RNA sequencing. Our implementation in Stan allows for easy extension to further sources of noise, technical or biological, while taking advantage of a state-of-the-art Hamiltonian Monte Carlo sampler. This approach allows us to accurately quantify uncertainty over parameter estimates and to perform Bayesian model selection to distinguish between modes of gene expression, or test for noise artefacts such as zero inflation.

While the telegraph model is known to fit experimentally observed mRNA distributions, it ignores cellspecific factors that provide an alternative explanation of measured mRNA counts. We show that accounting for extrinsic noise can significantly change the inferred dynamics of gene expression on a genome-wide level, as was also observed in [17, 18]. This affects burst size, frequency and duration, and our results suggest that some genes spend significantly more time in the on state than predicted, corroborating results in [17]. We uncovered little Bayesian evidence for zero inflation, consistent with [5], with the exception of selected genes when measuring single-allele mRNA counts.

Volume has a large effect on cellular gene expression levels and is an important factor of extrinsic noise [8], but this has to be separated from varying capture efficiencies between cells [19]. Separating these two is not straightforward in the absence of extrinsic controls, but we are positive that advances in scRNAseq technology will allow us to disentangle these effects more reliably in the future. Regardless of the type of noise considered, our results establish that genome-wide parameter inference can lead to significantly biased results when cell-to-cell variations are not taken into account. Our framework can be readily adapted to include various sources of technical and biological noise, such as cell cycle effects, when these can be reliably measured in scRNAseq experiments, and thus provides a robust and flexible starting point to analyse the mechanistic basis of genome-wide transcription.

## Acknowledgments

This work was supported by the EPSRC Centre for Doctoral Training in Data Science, funded by the UK Engineering and Physical Sciences Research Council (grant EP/L016427/1) and the University of Edinburgh. The author would like to thank Ramon Grima, Guido Sanguinetti and Augustinas Sukys for countless useful discussions and feedback.

## Data availability

Code for this publication is available at https://github.com/kaandocal/stan_rnaseq.

## References

[1] D. Jovic, X. Liang, H. Zeng, L. Lin, F. Xu, and Y. Luo. “Single-Cell RNA Sequencing Technologies and Applications: A Brief Overview”. Clin. Transl. Med. 12(3): e694 (2022).

[2] C. A. Vallejos, D. Risso, A. Scialdone, S. Dudoit, and J. C. Marioni. “Normalizing Single-Cell RNA Sequencing Data: Challenges and Opportunities”. Nat. Methods 14(6): 565–571 (2017).

[3] J. K. Kim and J. C. Marioni. “Inferring the Kinetics of Stochastic Gene Expression from Single-Cell RNA-sequencing Data”. Genome Biol. 14(1): R7 (2013).

[4] A. J. M. Larsson, P. Johnsson, M. Hagemann-Jensen, L. Hartmanis, O. R. Faridani, B. Reinius, Å. Segerstolpe, C. M. Rivera, B. Ren, and R. Sandberg. “Genomic Encoding of Transcriptional Burst Kinetics”. Nature 565(7738): 251 (2019).

[5] V. Svensson. “Droplet scRNA-seq is not Zero-Inflated”. Nat. Biotechnol. 38(2): 147–150 (2020).

[6] D. Grün, L. Kester, and A. van Oudenaarden. “Validation of Noise Models for Single-Cell Transcriptomics”. Nat. Methods 11(6): 637–640 (2014).

[7] A. M. Klein, L. Mazutis, I. Akartuna, N. Tallapragada, A. Veres, V. Li, L. Peshkin, D. A. Weitz, and M. W. Kirschner. “Droplet Barcoding for Single-Cell Transcriptomics Applied to Embryonic Stem Cells”. Cell 161(5): 1187–1201 (2015).

[8] R. Foreman and R. Wollman. “Mammalian Gene Expression Variability is Explained by Underlying Cell State”. Mol. Syst. Biol. 16(2): e9146 (2020).

[9] O. Padovan-Merhar, G. P. Nair, A. G. Biaesch, A. Mayer, S. Scarfone, S. W. Foley, A. R. Wu, L. S. Churchman, A. Singh, and A. Raj. “Single Mammalian Cells Compensate for Differences in Cellular Volume and DNA Copy Number through Independent Global Transcriptional Mechanisms”. Mol. Cell 58(2): 339–352 (2015).

[10] H. Kempe, A. Schwabe, F. Crémazy, P. J. Verschure, and F. J. Bruggeman. “The Volumes and Transcript Counts of Single Cells Reveal Concentration Homeostasis and Capture Biological Noise”. Mol. Biol. Cell 26(4): 797–804 (2015).

[11] J. Peccoud and B. Ycart. “Markovian Modeling of Gene-Product Synthesis”. Theor. Popul. Biol. 48(2): 222–234 (1995).

[12] P. V. Kharchenko, L. Silberstein, and D. T. Scadden. “Bayesian Approach to Single-Cell Differential Expression Analysis”. Nat. Methods 11(7): 740–742 (2014).

[13] C. A. Vallejos, J. C. Marioni, and S. Richardson. “BASiCS: Bayesian Analysis of Single-Cell Sequencing Data”. PLOS Comput. Biol. 11(6): e1004333 (2015).

[14] B. Carpenter, A. Gelman, M. D. Hoffman, D. Lee, B. Goodrich, M. Betancourt, M. A. Brubaker, J. Guo, P. Li, and A. Riddell. “Stan: A Probabilistic Programming Language”. J. Stat. Softw. 76: 1 (2017).

[15] W. Tang, F. Bertaux, P. Thomas, C. Stefanelli, M. Saint, S. Marguerat, and V. Shahrezaei. “bayNorm: Bayesian Gene Expression Recovery, Imputation and Normalization for Single-Cell RNA-sequencing Data”. Bioinformatics 36(4): 1174–1181 (2020).

[16] B. Pijuan-Sala, N. K. Wilson, J. Xia, X. Hou, R. L. Hannah, S. Kinston, F. J. Calero-Nieto, O. Poirion, S. Preissl, F. Liu, and B. Göttgens. “Single-Cell Chromatin Accessibility Maps Reveal Regulatory Programs Driving Early Mouse Organogenesis”. Nat. Cell Biol. 22(4): 487–497 (2020).

[17] X. Fu, H. P. Patel, S. Coppola, L. Xu, Z. Cao, T. L. Lenstra, and R. Grima. “Quantifying How Post-Transcriptional Noise and Gene Copy Number Variation Bias Transcriptional Parameter Inference from mRNA Distributions”. eLife 11: e82493 (2022).

[18] R. Grima and P.-M. Esmenjaud. Quantifying and Correcting Bias in Transcriptional Parameter Inference from Single-Cell Data. 2023. biorxiv: 10.1101/2023.06.19.545536. preprint.

[19] W. Tang, A. C. S. Jørgensen, S. Marguerat, P. Thomas, and V. Shahrezaei. “Modelling Capture Efficiency of Single-Cell RNA-sequencing Data Improves Inference of Transcriptome-Wide Burst Kinetics”. Bioinformatics 39(7): btad395 (2023).

[20] V. A. Herzog, B. Reichholf, T. Neumann, P. Rescheneder, P. Bhat, T. R. Burkard, W. Wlotzka, A. von Haeseler, J. Zuber, and S. L. Ameres. “Thiol-Linked Alkylation of RNA to Assess Expression Dynamics”. Nat. Methods 14(12): 1198–1204 (2017).

[21] L. Ham, R. D. Brackston, and M. P. H. Stumpf. “Extrinsic Noise and Heavy-Tailed Laws in Gene Expression”. Phys. Rev. Lett. 124(10): 108101 (2020).

[22] C. Ziegenhain, G.-J. Hendriks, M. Hagemann-Jensen, and R. Sandberg. “Molecular Spikes: A Gold Standard for Single-Cell RNA Counting”. Nat. Methods 19(5): 560–566 (2022).

[23] R. Jiang, T. Sun, D. Song, and J. J. Li. “Statistics or Biology: The Zero-Inflation Controversy about scRNA-seq Data”. Genome Biology 23(1): 31 (2022).

[24] Y. Jiang, N. R. Zhang, and M. Li. “SCALE: Modeling Allele-Specific Gene Expression by Single-Cell RNA Sequencing”. Genome Biol. 18(1): 74 (2017).

[25] C. J. Hsiao, P. Tung, J. D. Blischak, J. E. Burnett, K. A. Barr, K. K. Dey, M. Stephens, and Y. Gilad. “Characterizing and Inferring Quantitative Cell Cycle Phase in Single-Cell RNA-seq Data Analysis”. Genome Res. 30(4): 611–621 (2020).

[26] A. Riba, A. Oravecz, M. Durik, S. Jiménez, V. Alunni, M. Cerciat, M. Jung, C. Keime, W. M. Keyes, and N. Molina. “Cell Cycle Gene Regulation Dynamics Revealed by RNA Velocity and Deep-Learning”. Nat. Commun. 13(1): 2865 (2022).

[27] D. M. Higdon. “Auxiliary Variable Methods for Markov Chain Monte Carlo with Applications”. J. Am. Stat. Assoc. 93(442): 585–595 (1998).

[28] A. R. Stinchcombe, C. S. Peskin, and D. Tranchina. “Population Density Approach for Discrete mRNA Distributions in Generalized Switching Models for Stochastic Gene Expression”. Phys. Rev. E 85(6): 061919 (2012).

[29] Y. Zhou, J. S. Hagood, and J. E. Murphy-Ullrich. “Thy-1 Expression Regulates the Ability of Rat Lung Fibroblasts to Activate Transforming Growth Factor-Beta in Response to Fibrogenic Stimuli”. Am. J. Pathol. 165(2): 659–669 (2004).

[30] R. D. Morey, J. N. Rouder, M. S. Pratte, and P. L. Speckman. “Using MCMC Chain Outputs to Efficiently Estimate Bayes Factors”. J. Math. Psychol. 55(5): 368–378 (2011).

