## Supplementary Information for "Incorporating extrinsic noise into mechanistic modelling of single-cell transcriptomics"

### A The telegraph and extrinsic noise models

To model mRNA counts we used the Beta-Poisson model [28], an alternative description of the telegraph model at steady state:

$$z \sim \mathrm{Beta}(\sigma_{\mathrm{on}}, \sigma_{\mathrm{off}}), \qquad n \mid z \sim \mathrm{Poi}(\rho z). \tag{A1}$$

This model describes gene expression from a single allele. We also considered the pooled telegraph model to describe expression from two independent alleles:

$$z_1, z_2 \sim \mathrm{Beta}(\sigma_{\mathrm{on}}, \sigma_{\mathrm{off}}), \qquad n \mid z \sim \mathrm{Poi}(\rho\,(z_1 + z_2)). \tag{A2}$$

This can be extended to account for gene duplication after the S-stage, depending on the cells studied.
An alternative for pooled data is to use the classical telegraph model; in this case the average transcription rate $\rho$ per allele can be estimated by halving the empirical transcription rate, and similarly for $\mu$ and $b$.

We modelled zero-inflation by adding an artificial peak at 0 to the raw mRNA distribution, resulting in a zero-inflated distribution of the form

$$P_{\mathrm{zi}}(n) = p_0\,\delta(n) + (1 - p_0)\,P(n), \tag{A3}$$

where $P(n)$ is the distribution of the original model and $p_0 \in [0, 1]$ describes the fraction of zero reads due to dropout (as opposed to the total fraction of zero reads). Testing for the existence of zero inflation amounts to checking whether $p_0 = 0$. We emphasise that this model of zero inflation does not include increased zero counts due to binomial thinning.

For the extrinsic noise model we added the cell-specific parameters $V_c > 0$, subject to the normalisation condition

$$\frac{1}{N} \sum_c V_c = 1, \tag{A4}$$

and defined the cell-specific transcription rate $\rho_c$ following (5).

To obtain the mechanistic parameters of the telegraph model starting from our reparametrisation (4) we use

$$\rho = \frac{b}{d}, \qquad \sigma_{\mathrm{on}} = \frac{\mu}{b - \mu d}, \qquad \sigma_{\mathrm{off}} = d^{-1}, \tag{A5}$$

which is valid under our assumption that $b > \mu d$. This is the natural requirement that the average number of mRNA produced cannot exceed the amount of mRNA produced per burst, divided by the average length of a burst.

### B Model priors

We used the following priors for the zero-inflated telegraph model:

$$\mu \sim \mathrm{Exp}(20^{-1}), \qquad d \sim \mathrm{Exp}(1), \tag{B1}$$

$$b \sim d\mu + \mathrm{Exp}(20^{-1}), \qquad p_0 \sim \mathrm{U}(0, 1). \tag{B2}$$

For the extrinsic noise model we chose a uniform prior on the volume factors $V_c$, conditioned on (A4). This is equivalent to a rescaled Dirichlet distribution with concentration parameter 1. The prior on the parameter $\alpha$ is given by

$$\alpha \sim \mathrm{U}(0, 1). \tag{B3}$$

### C MCMC sampling

We used Stan [14] to sample the posterior over the parameters for each of the considered models. For all experiments we ran 8 independent chains to check for convergence of the posterior by comparing the variance between chains with that within chains, and by visual inspection. Each chain was run for 1k warmup iterations, followed by 30k sampling steps that were thinned by a factor of 10, resulting in 3k posterior samples per chain.

#### D Bayesian model selection

The Bayes factor compares the evidence for two models that are nested in our case: $\mathcal{M}_0 \subseteq \mathcal{M}$, where $\mathcal{M}_0$ is defined by $\phi = 0$ for some parameter $\phi$; this could be the amount of zero inflation $p_0$, or the mean burst duration $d$. Assuming that the prior for $\mathcal{M}_0$ is obtained by restricting the prior $\mathcal{M}$ to $\phi = 0$, we can write the Bayes factor as the Savage-Dickey ratio

$$\mathrm{BF} = \frac{p(\mathcal{D} \mid \mathcal{M}_0)}{p(\mathcal{D} \mid \mathcal{M})} = \frac{p(\phi = 0 \mid \mathcal{D})}{p(\phi = 0)} \tag{D1}$$

This ratio can be computed from the posterior for $\mathcal{M}$ by estimating the density at $\phi = 0$, or using the method in [30]. We instead computed the approximation

$$\widehat{\mathrm{BF}} = \frac{p(\phi < \epsilon \mid \mathcal{D})}{p(\phi < \epsilon)}, \tag{D2}$$

for a tolerance parameter $\epsilon = 0.02$. This ratio can be estimated by counting the fraction of posterior samples below the threshold. Our results did not significantly change when $\epsilon$ was varied between 0.01 and 0.03.

##### The bursty limit

Here we compared the evidence for the telegraph model $\mathcal{M}$ with the bursty limit $\mathcal{M}_{\mathrm{bur}}$, defined by $d \approx 0$ in the telegraph model, via the Bayes factor

$$\mathrm{BF}_{\mathrm{bur}} = \frac{p(\mathcal{D} \mid \mathcal{M}_{\mathrm{bur}})}{p(\mathcal{D} \mid \mathcal{M})} \tag{D3}$$

for each gene. In practice we computed the approximation (D2); this does not significantly bias our findings since on durations of less than 0.02, normalised by $\delta$, are practically indistinguishable from the bursty model at steady state.

The Bayes factor (D3) depends on the prior chosen, and particularly on the prior over $d$. The latter, described in SI B, is a reasonable choice given the expected scale of $d$ across genes. Since $d$ is normalised by the mean RNA lifetime, values of $d$ much larger than 1 imply that a significant fraction of mRNA is degraded before the gene turns off, which is not expected behaviour for most genes.

##### Zero inflation

To test for the existence of zero inflation we compared the evidence for the original model $\mathcal{M}$ with that for the zero-inflated model $\mathcal{M}_{\mathrm{zi}}$ and computed the Bayes factor

$$\mathrm{BF}_{\mathrm{zi}} = \frac{p(\mathcal{D} \mid \mathcal{M}_{\mathrm{zi}})}{p(\mathcal{D} \mid \mathcal{M})} \tag{D4}$$

for each gene. Note that here the original model $\mathcal{M}$ is a special case of the zero-inflated model $\mathcal{M}_{\mathrm{zi}}$ with $p_0 = 0$, and (D4) is the reciprocal Savage-Dickey ratio. The uniform prior over $p_0$ represents a complete lack of knowledge about the extent of zero-inflation. As for the bursty case we used (D2) to estimate the evidence for zero inflation.

The excess 0 ratio in (Fig. 2**d)** and Fig. 2**d)** was obtained by comparing the estimated amount of zero inflation, measured by the inferred value of $p_0$, with the expected number of zero reads. The latter was coarsely estimated by assuming a roughly uniform distribution on mRNA numbers; in this case the probability of a zero read is approximately $1/2\mu$. We therefore defined the excess 0 ratio as

$$2 p_0 \mu. \tag{D5}$$

### E Intrinsic and extrinsic variance

For a model of the form

$$n \mid V, \lambda \sim \mathrm{Poi}(\lambda(\alpha V + 1 - \alpha)), \tag{E1}$$

where $\lambda > 0$ represents an arbitrary latent variable, we have the following variance decomposition:

$$\sigma^2 =: \mathrm{Var}(n) = \mu(1 + \mathrm{FF}_\lambda) + \mu^2\alpha^2\,\mathrm{CV}_V(1 + \mathrm{CV}_\lambda), \tag{E2}$$

cf. [18]. This allows us to compute the extrinsic noise as

$$\sigma_e^2 = \mu^2\alpha^2\,\mathrm{CV}_V(1 + \mathrm{CV}_\lambda). \tag{E3}$$

For the telegraph model, the latent variable $\lambda$ follows a rescaled Beta distribution:

$$\lambda \sim \rho \cdot \mathrm{Beta}(\sigma_{\mathrm{on}}, \sigma_{\mathrm{off}}). \tag{E4}$$

Adjusting the inferred parameters for the capture effiency $\beta$ requires dividing the variable $\lambda$ by $\beta$. Doing this we obtain

$$\sigma^2(\beta) = \frac{\mu}{\beta}\left(1 + \frac{\mathrm{FF}_\lambda}{\beta}\right) + \frac{\mu^2}{\beta^2}\alpha^2\,\mathrm{CV}_V(1 + \mathrm{CV}_\lambda). \tag{E5}$$

As long as $\mathrm{FF}_\lambda \gg 1$ (which excludes Poisson statistics), the intrinsic and extrinsic noise terms scale roughly with $\beta^{-2}$ and the variance decomposition is robust to different values of $\beta$.

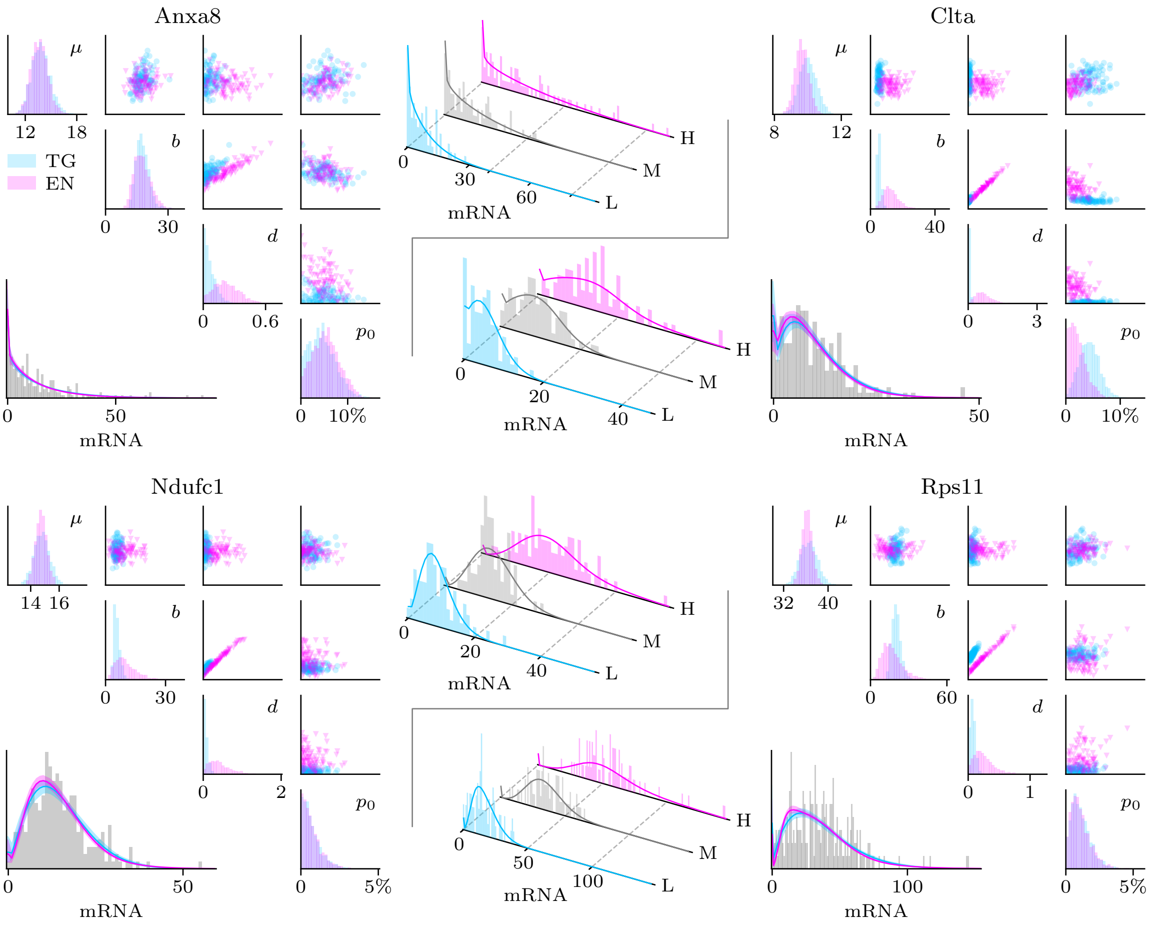


**Figure S1:** Comparison of inference results for the telegraph and extrinsic noise models, showing 4 selected genes (CAST allele). See Fig. 1 for explanation.

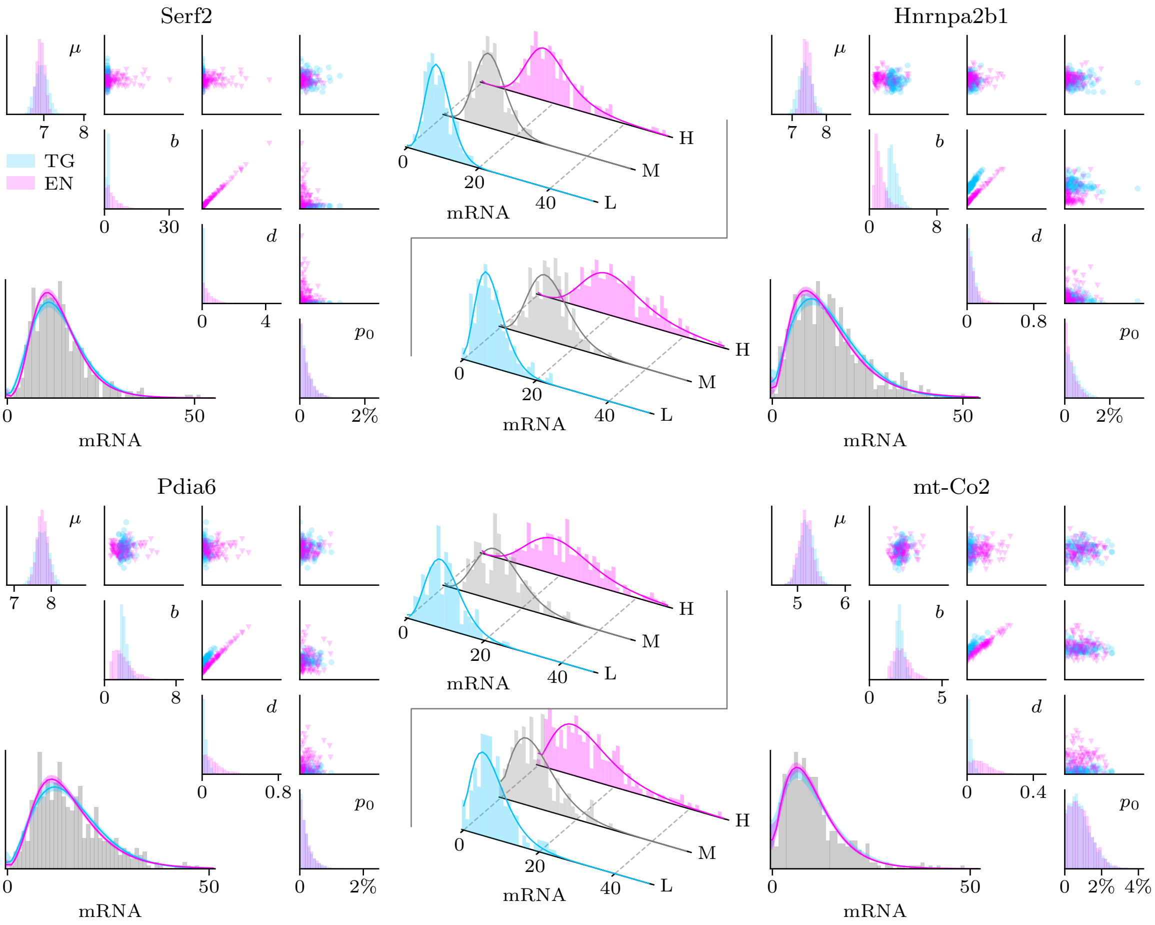


**Figure S2:** Comparison of inference results for the telegraph and extrinsic noise models, showing 4 selected genes (mESC data, replicate 1). See Fig. 1 for explanation. The parameters $\mu$ and $b$ are normalised for one allele as described in SI A.

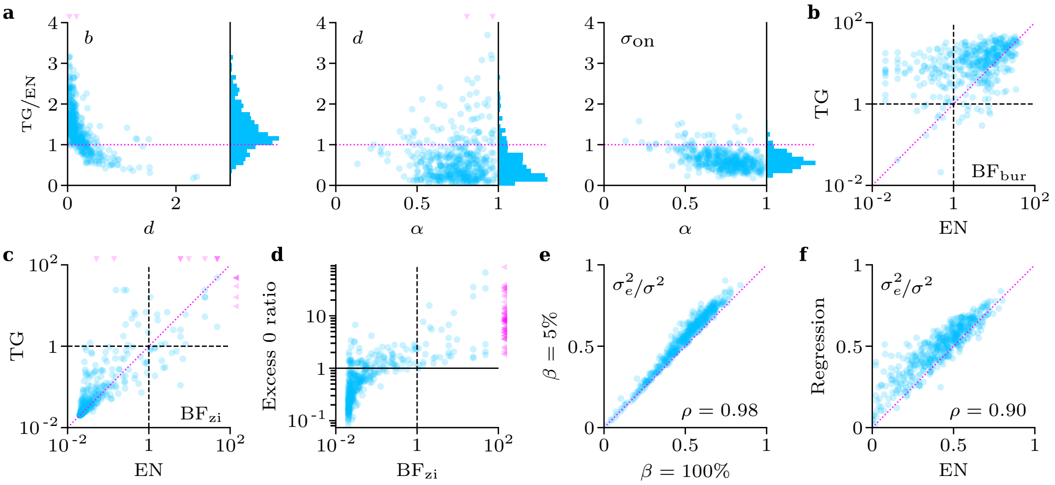


**Figure S3:** Analysis of the single-allele mouse fibroblast dataset (CAST allele) using the telegraph (TG) and extrinsic noise (EN) models. See Fig. 2 for explanation.

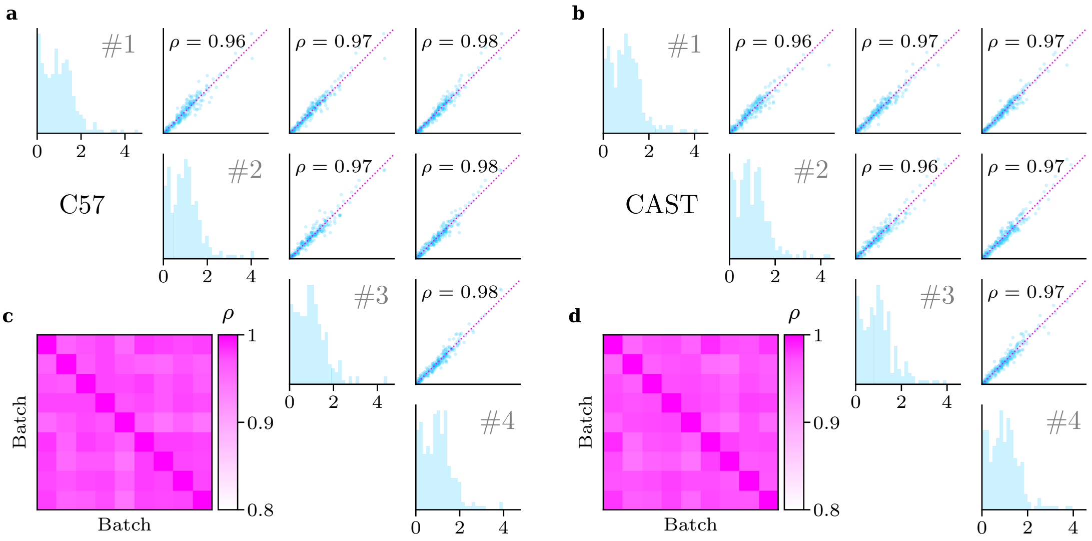


**Figure S4:** Cell-specific volume factors $V_c$ are consistently estimated between batches (fibroblast data, both alleles). **a, b)** The diagonals contain histograms of $V_c$ across cells for each batch batches (selection only). Pairwise comparisons of estimates between batches are plotted above the diagonal. **c, d)** Heatmap of the Pearson correlation $\rho$ between batches.

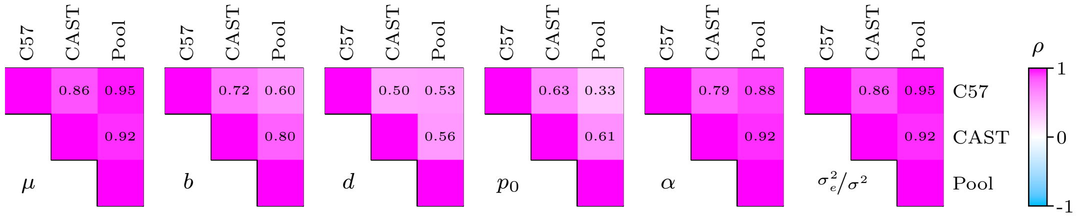


**Figure S5:** Comparison of parameters estimated for the fibroblast dataset on a single-allele basis and from pooled data.

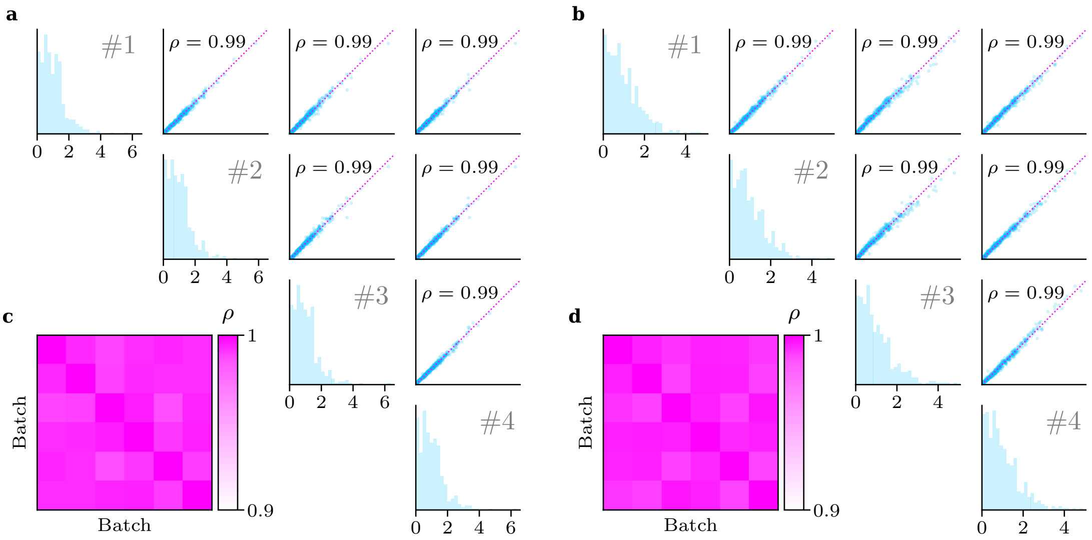


**Figure S6:** Cell-specific volume factors $V_c$ are consistently estimated between batches (mESC data, replicates 1 and 2). See Fig. S4 for explanation.

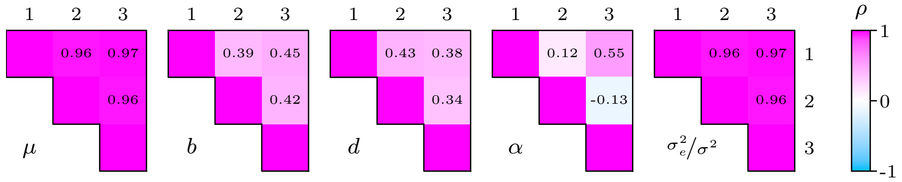


**Figure S7:** Comparison of parameters estimated for the mESC dataset across three batches, using the pooled telegraph model.
